# High species turnover shapes anuran community composition in ponds along an urban-rural gradient

**DOI:** 10.1101/2020.09.01.276378

**Authors:** Carolina Cunha Ganci, Diogo B. Provete, Thomas Püttker, David Lindenmayer, Mauricio Almeida-Gomes

## Abstract

The rapid expansion of urban areas in which natural and semi-natural areas are replaced by human infrastructure, such as buildings and streets, is a major threat to biodiversity worldwide. However, little is known about how the structure of biotic communities is affected by urbanization in the tropics. Here, we tested the effect of land use types in urban and peri-urban areas on frog species richness and community composition in central Brazil. We selected 20 ponds differing in size and surrounding levels of urbanization as well as natural forest cover. We then used a Poisson GLM and a distance-based Redundancy Analysis (db-RDA) to relate species richness and community composition, respectively, to environmental variables. Variation in species richness was best explained by pond size (positive effect) and amount of urbanization (negative effect) in the surrounding 500 m. Community composition was mainly driven by species turnover than by nestedness, with db-RDA showing that turnover was explained primarily by urban infrastructure and forest cover. Our results indicate that urbanization negatively influences species richness. Moreover, as the amount of urbanization increased, several species were replaced by others taxa that appear better adapted to urban environments. Our results indicate that maintaining large ponds with surrounding native vegetation in urban environments might be an effective strategy for conserving frog communities.

## 1. Introduction

Urbanization is a key threat to biodiversity, causing extirpation of populations and species worldwide (Miller and Hobbs 2002, McKinney 2008, Batáry et al. 2018). Currently, more than 50% of the world’s human population lives in urban areas and it is estimated that this number will increase to approximately 65% by 2030 (Grimm et al. 2008, United Nations 2018). Urbanization can drastically alter the chemical, physical, and ecological condition of ecosystems, with several studies highlighting the negative effects of urbanization on species dispersal (Findlay et al. 2001, Rubbo and Kiesecker 2005), ecological interactions (Fischer et al. 2012), and community structure (Christie et al. 2010, Zhang et al. 2016, Gagné and Fahrig 2007, McDonnell et al. 1997, Kinzig and Grove 2001, Johnson et al. 2013). For example, in an investigation of bird communities along a gradient of urban influence, Dale (2018) found that urbanization led to a predominance of species with generalist diets and nesting habits.

Differences in composition among communities (beta-diversity, Whittaker 1960) may reflect two distinct ecological processes: species loss (nestedness), in which the composition of species-poor assemblages represents a subset of richer sites (Bender et al. 2017), or replacement (turnover) of species, in which sets of species are replaced by others (Baselga 2010, Anderson et al. 2011, Leibold and Chase 2018). Urbanization has been shown to affect species richness leading to nested communities in cities compared to less urbanized areas (González-Oreja et al. 2012, Ficetola and De Bernardi 2004). For example, Fernández-Juricic (2002) showed that bird species exhibited a nested pattern in areas subject to high levels of human influence. Similarly, taxonomic composition of ant communities in most urbanized locations were a subset of the least disturbed locations (Slipinski et al. 2012), and dragonfly beta diversity in urban stormwater ponds was explained mostly by differences in richness rather than by turnover (Johansonn et al. 2019).

Amphibians are the most threatened group of vertebrates globally, with approximately 40% of its species classified in at least one threat category of the IUCN (IUCN 2020). Nearly 1,000 amphibian species are classified as threatened with extinction because of land use changes for urban development (Maxwell et al. 2016), likely due to the fact that most amphibian species exhibit complex life cycles (Wells 2007), and urbanization can seriously affect both aquatic and terrestrial environments (Zhang et al. 2016). For example, Knutson et al. (1999) showed that urban development affected both tadpole and adult stages, by modifying terrestrial environments as well contaminating water bodies. Similarly, major changes in landscape composition through urbanization severely impacted anuran populations, affecting richness, abundance, and occurrence of several species (Ficetola et al. 2011, Youngquist et al. 2017). In the few studies investigating specific components of urban landscapes, pond size has generally proved to be a strong predictor of species richness and abundance (Dickman 1987, Parris 2006, Gagné and Fahrig 2007), although its effect on community composition is unclear.

Despite of anuran diversity being strongly affected by urbanization, amphibians are among the least studied vertebrate groups in urban landscapes worldwide (McDonnell and Hahs 2008). Although the number of studies on amphibians in urban areas has grown, information is still lacking to effectively guide amphibian conservation in most of the world’s urbanized landscapes (but see Parris et al. 2018), especially in the highly diverse tropics. Here, we tested the effect of landscape composition on anuran communities in ponds along an urban-rural gradient. We hypothesized that anuran species richness in cities is positively related to habitat area and negatively related to urbanization. We also investigated which process (nestedness or turnover) was responsible for differences in anuran community composition along an urban-rural gradient. We hypothesized that decreasing habitat area and increasing urbanization would lead to the local extinction of sensitive species without replacement of species. To the best of our knowledge, this is the first study testing the contribution of separate components of beta diversity on changes in anuran community composition across an urban-rural gradient.

## 2. Methods

### 2.1 Study site

We conducted this study in Campo Grande city (20° 28’ 13”S, 54° 37’ 25”W), Mato Grosso do Sul state, central Brazil. This region was originally covered by savanna vegetation, with scattered trees and shrubs mixed with forest formations (known as Cerrado; Ribeiro and Walter, 1998). Currently only ~ 21% of the original native cover persists, while the remaining 79% of the municipally consists primarily of pastures, agricultural fields, and urban areas (PLANURB 2019). The climate is characterized by a dry season (between April to September) and a rainy season (October to March), with an annual average rainfall of 1,530 mm, and an average annual temperature ranging from 18 °C to 29 °C (Ferreira et al. 2017). The urban area of Campo Grande covers 154.4 km^2^, with ~ 900,000 inhabitants, and a mean population density of 104 inhabitants/km^2^ (Ferreira et al. 2017).

### 2.2 Sampling design

We georeferenced and visited 156 ponds in urban and peri-urban areas, and selected 20 ponds for survey (Fig 1). We based pond selection on a careful systematic procedure. First, we included only ponds that were a minimum distance of 1 km apart (1.1 to 26.7 km), to maximize independence of sampling units (Semlisch and Bodie 2003). We tested for independence by completing a Mantel correlogram and found no evidence of spatial autocorrelation in our data (Fig. S1). Second, we selected ponds that maximized variation of explanatory variables (i.e. amount of surrounding native forest cover, degree of urbanization, and pond size; Table S1). Third, we selected ponds to encompass a broad range of pond sizes (109 to 3,080 m^2^; Table S1).

**Fig. 1.**
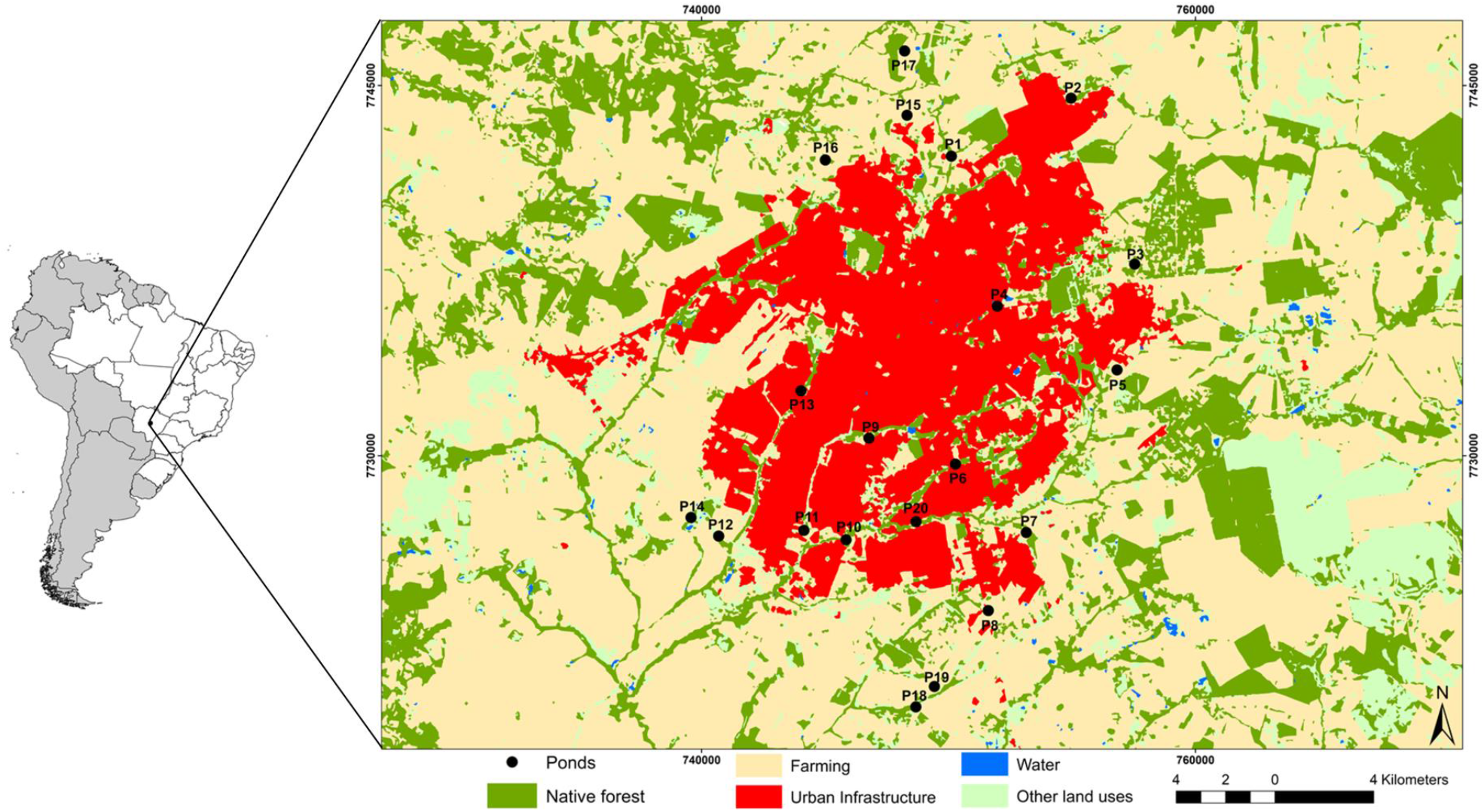
Study area showing the location of the 20 ponds sampled in urban and peri-urban areas in Campo Grande, Mato Grosso do Sul, central Brazil. Land use classification was extracted from the MapBiomas database for 2018.

### 2.3 Frog sampling

We estimated the abundance of frogs through visual encounter surveys (Crump and Scott Jr 1994) at night (19:00 h to 00:00 h), using headlamps. This is considered the most efficient method to detect anurans (Doan 2003, Flint and Harris 2005, Grover 2006). We also used acoustic identification because some anuran species have low detectability due to their small body size, rarity, or their ability to camouflage among the leaf litter or underground (Sá et al. 2018, Lima et al. 2019). However, while this alternative method allowed for the detection of species with predominantly subterranean habits (which otherwise would have been overlooked), only the abundances of two comparatively rare species (*Elachistocleis helianneae* and *Elachistocleis bicolor*) were estimated by acoustic identification. We therefore assumed there was low impact of the two different methods on abundance estimates.

We visited each pond three times on different days between October of 2018 and December 2018, with each visit lasting 30 minutes and performed by three people. This gave a sampling effort of 4.5 h per pond and 90 h in total. During the surveys, researchers walked the entire perimeter of the pond more than once, looking for individuals at the edge of, and inside, ponds. We identified each individual to species level and then marked it with Visible Implant Elastomer (VIE) Tags (Northwest Marine Technology, Inc.), to avoid recounting the same individual. This is an alternative to the traditional toe-clipping method, but does not harm animals (Davis and Ovaska 2001). Mean sampling coverage (proportion of observed richness relative to richness estimated by Chao 1) for all ponds was 0.9 (range: 0.41 to 1), indicating that sampling effort was sufficient to estimate species presence in most local communities (Table S2).

### 2.4 Landscape metrics

We obtained information on land use types based on intermediate resolution images (30 m) provided by the MapBiomas platform (http://mapbiomas.org; collection 4.0), for 2018. We calculated the percentages of land use classes around ponds using ARCGIS 10.6.1. For each pond, we measured pond size (m^2^; through polygon tool, available in Google Earth), and the percentage of native forest as well as urban infrastructure in buffer sizes of 500 and 1,000 m radius. We defined native forest using the classification “Natural Forest” provided by MapBiomas, including forest and savanna formations. Urban infrastructure included all non-native land uses, like roads and buildings. We selected the two buffer sizes based on previous studies (Semlitsch and Bodie 2003, Almeida-Gomes et al. 2016). We did not add smaller scales due to very low variation in land use coverage (Table S1), thereby precluding analyses. Likewise, we excluded buffers at larger scales (> 1,000 m) due to high levels of overlap with nearby buffers, thereby potentially leading to pseudoreplication. The percentage of native forest cover varied from 0.3 to 36.1% in a 500 m buffer and 1.8 to 47.3% in a 1,000 m buffer. Urban infrastructure varied from 0 to 89.4% in a 500 m buffer and 0 to 84.5% in a 1,000 m buffer (Table S1).

### 2.5 Data analysis

We standardized all five predictor variables to zero mean and unit variation prior to analysis. We used the variance inflation factor (VIF) in the usdm package (Naimi et al. 2014) to check for multicollinearity. Our final dataset only included uncorrelated variables (VIF < 10): pond size, native forest in 500 m and 1,000 m buffer, and urban infrastructure in 500 m buffer.

#### 2.5.1 Species richness

To test the effects of forest cover (native forest), urbanization (urban infrastructure), and pond size on amphibian richness, we used generalized linear models (GLMs) with Poisson distributed residuals. We used the sjstats package (Lüdecke 2019) to check for overdispersion and conduct residual diagnosis. All models had normally and homogeneously distributed residuals. We found no evidence of overdispersion.

We built and fit 12 candidate models to test alternative hypotheses about the importance of groups of variables in explaining amphibian species richness: three models to test the effect of forest cover, urbanization, and pond size at two spatial scales, one model for each spatial scale to test the effect of urbanization and forest cover, and one model for each spatial scale to test the combined effect of pond size, urbanization, and forest cover. We used Akaike Information Criterion corrected for small samples (AICc) and Akaike weights to rank models, considering as plausible all models with ΔAICc ≤ 2. We conducted analyses in the MuMIn package (Barton 2018).

#### 2.5.2 Beta diversity partitioning

We decomposed total beta diversity into two components: turnover and nestedness, following Baselga (2010). We used presence-absence and abundance data to calculate dissimilarity and its two components using Jaccard and Bray-Curtis indices, respectively. We conducted our analyses in the betapart R package (Baselga et al. 2018) and the three dissimilarity matrices were summarized using Principal Coordinates Analysis. After these analyses, we performed a distance-based Redundancy Analysis (db-RDA) to test which landscape variables best explained the nestedness or turnover distance matrices. We conducted the analysis in the vegan package (Oksanen et al. 2019). db-RDA exhibited a high level of performance, with a negligible false-negative rate (Jupke and Schäfer 2020), and was an adequate constrained ordination method to relate distance matrices to predictor variables. As we did not detect a high degree of spatial autocorrelation in the community composition (Fig. S1), we did not include any spatial predictors (e.g., Moran Eigenvector Maps) in our model. We completed all analyses in R v. 3.6.0 (R Core Team 2019).

## 3. Results

We found 835 individuals of 20 anuran species from five families (Table S3). The most speciose families were Hylidae and Leptodactylidae (nine and seven species, respectively). The most abundant species were *Dendropsophus nanus*, with 266 individuals, which represented 31.8% of the total sampled individuals and *Scinax fuscomarginatus* (87; 10.4%). Frog richness in ponds ranged from one to 13 species and abundance from six to 139 individuals.

We found a single best-fit model to explain frog species richness (w_i_ = 0.608; R^2^ = 0.804), which contained pond size and percentage of urban infrastructure in a 500 m buffer (Table 1). Our analyses revealed that pond size (Coefficient ± SE: 0.22 ± 0.07) and urban infrastructure (- 0.31 ± 0.11) had a positive and negative effect on species richness, respectively (Fig. 2).

**Table 1.**
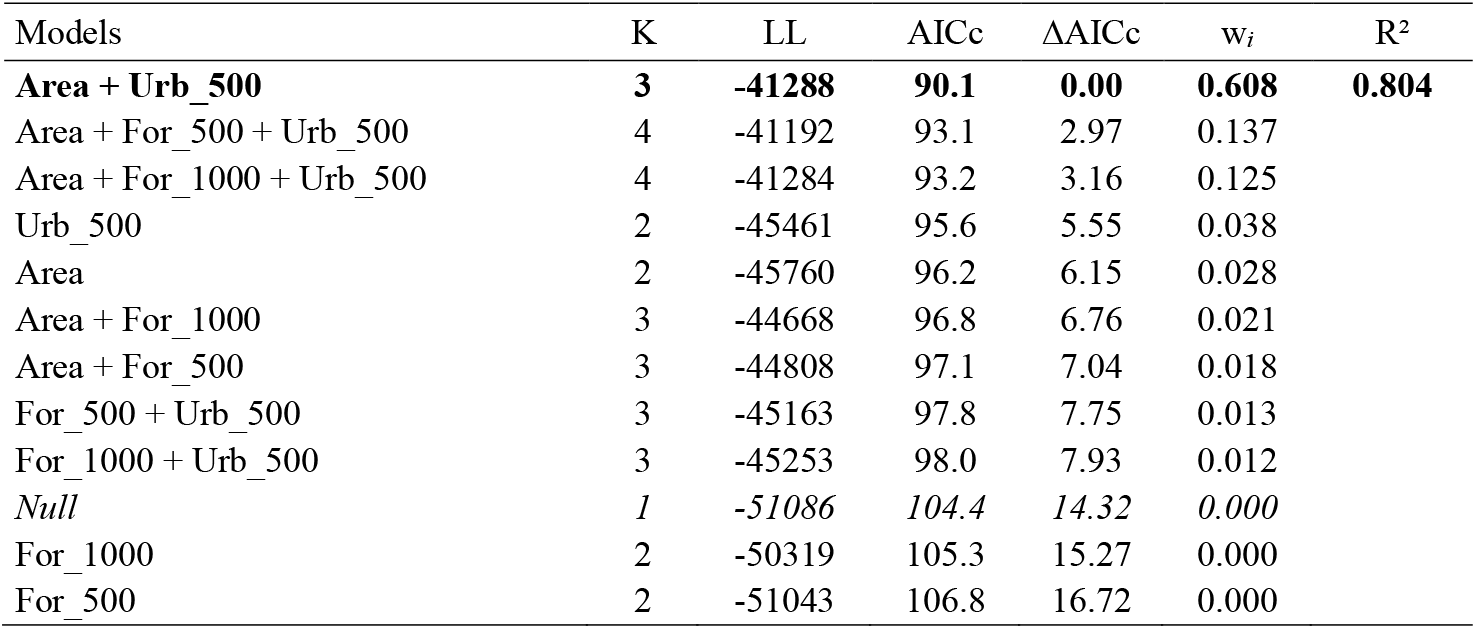
Results of model selection based on Akaike Information Criterion (AICc) explaining anuran richness. We highlight the best model and the null model in bold and italics, respectively. Area – pond size in m^2^; For_500 – percentage of native forest in a buffer of 500 m; For_1000 – percentage of native forest in a buffer of 1000 m; Urb_500 – percentage of urban infrastructure in a buffer of 500 m; LL – model log likelihood; w_i_ - Akaike weights; R^2^ - Nagelkerke’s R^2^ (1991).

**Fig. 2.**
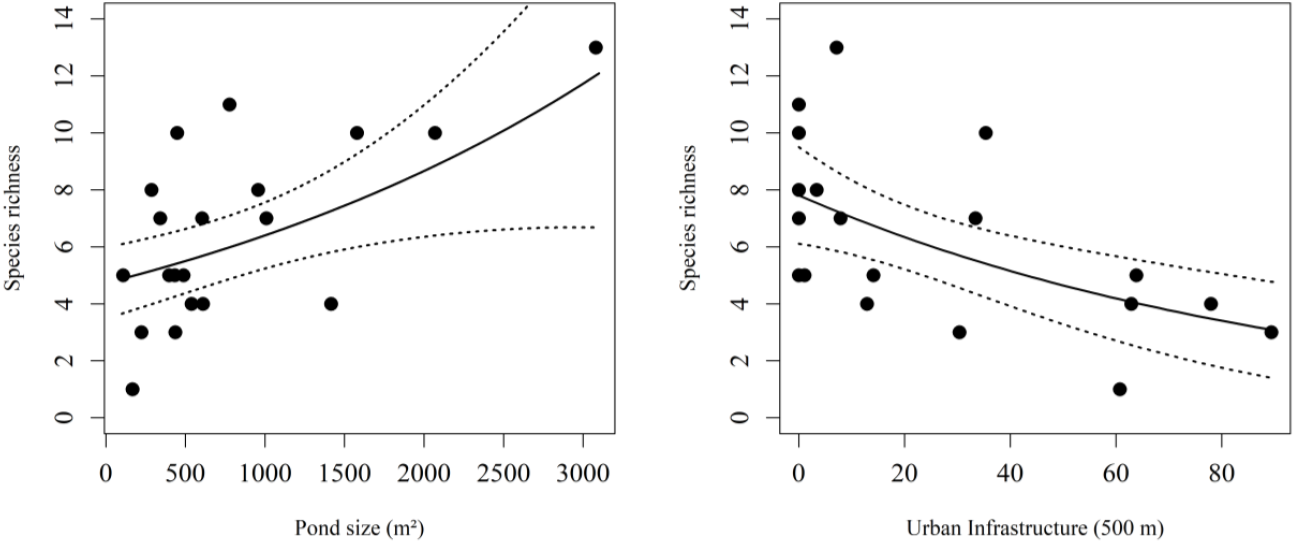
Effects of landscape metrics on anuran species richness. A) Positive relationship between pond size (m^2^) and species richness, after controlling for urban infrastructure in a 500-m buffer. B) Negative effect of percentage of urban infrastructure in a 500-m buffer (%) on species richness, after controlling for pond size. Solid lines represent the estimated relationship among variables, and dotted lines are 95% confident intervals.

The composition of amphibian communities in ponds differed considerably, ponds with high urban infrastructure were distinguished from those in less urbanized areas in the ordination diagram (Fig. 3). This separation was mostly explained by turnover (92.7%) rather than nestedness (7.3%). Notably, ponds with similar percentage of urbanization were more similar in composition indicating biotic homogenization along the gradient (Fig. 3). The overall results did not change if we used data on species abundance (Fig. S2).

**Fig. 3.**
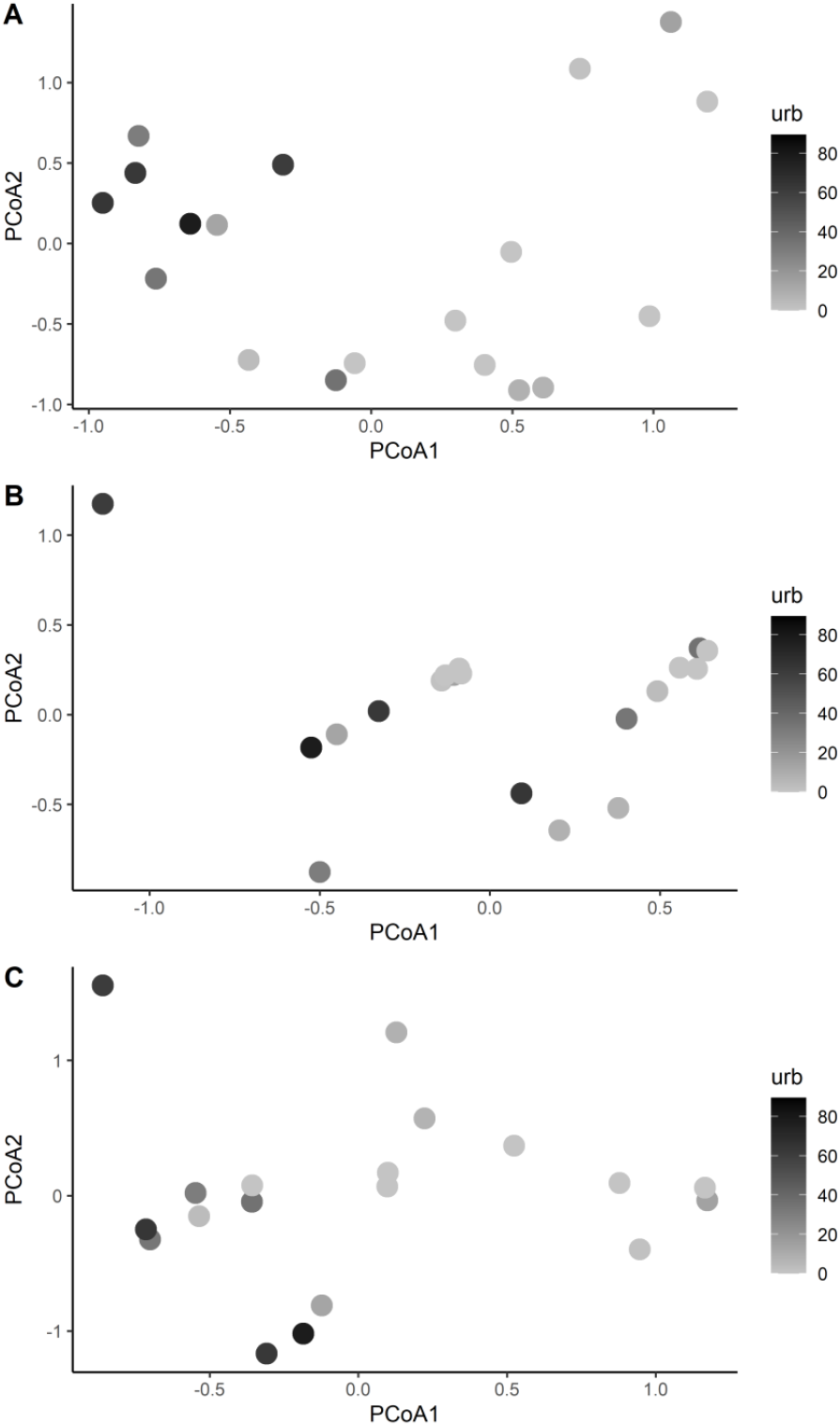
Ordination diagram showing the Principal Coordinate Analysis for the total (A), nestedness (B), and turnover (C) pair-wise dissimilarity matrix between sites, considering presence-absence data. Symbols are shaded according to the amount of urban infrastructure in a buffer of 500 m surrounding each pond (Urb_500).

Our application of db-RDA showed (r^2^ = 0.34, P = 0.02) that the variables that best explained turnover were the percentage of urban infrastructure in a 500 m buffer (F_3,16_ = 4.32, *P* = 0.002) and native forest in a buffer of 500 m (F_3,16_ = 2.47, *P* = 0.035; Fig. S3). The nestedness component (r^2^ = 0.72, P = 0.03) was best explained by pond size (F_3,16_ = 2.91, *P* = 0.036; Fig. S4), and the total beta diversity (r^2^ = 0.23, P = 0.02) was best explained by pond size (F_3,16_ = 2.26, P = 0.021; Fig. S4), percentage of native forest in a 500 m buffer (F_3,16_ = 2.20, *P* = 0.031; Fig. S4) and, percentage of urbanization in a 500 m buffer (F3,16 = 4.32, *P* = 0.001; Fig. S4).

The proportion of black (presence) and gray (absence) blocks in the diagram ordered by the urbanization gradient (Fig. 4) added to the evidence that the community was structured by turnover. When the percentage of urbanization increased, some species were replaced by others. For example, in a comparison of pond 12 (0% urbanization) and pond 2 (35% urbanization), we found that both supported the same number of species (n = 10), but the composition was different, with *Leptodactylus fuscus, Physalaemus nattereri*, and *Elachistocleis helianneae* absent from pond 2, but present in pond 12, and the addition of *Boana punctata, Leptodactylus labyrinthicus*, and *Dendropsophus minutus* to pond 2.

**Fig. 4.**
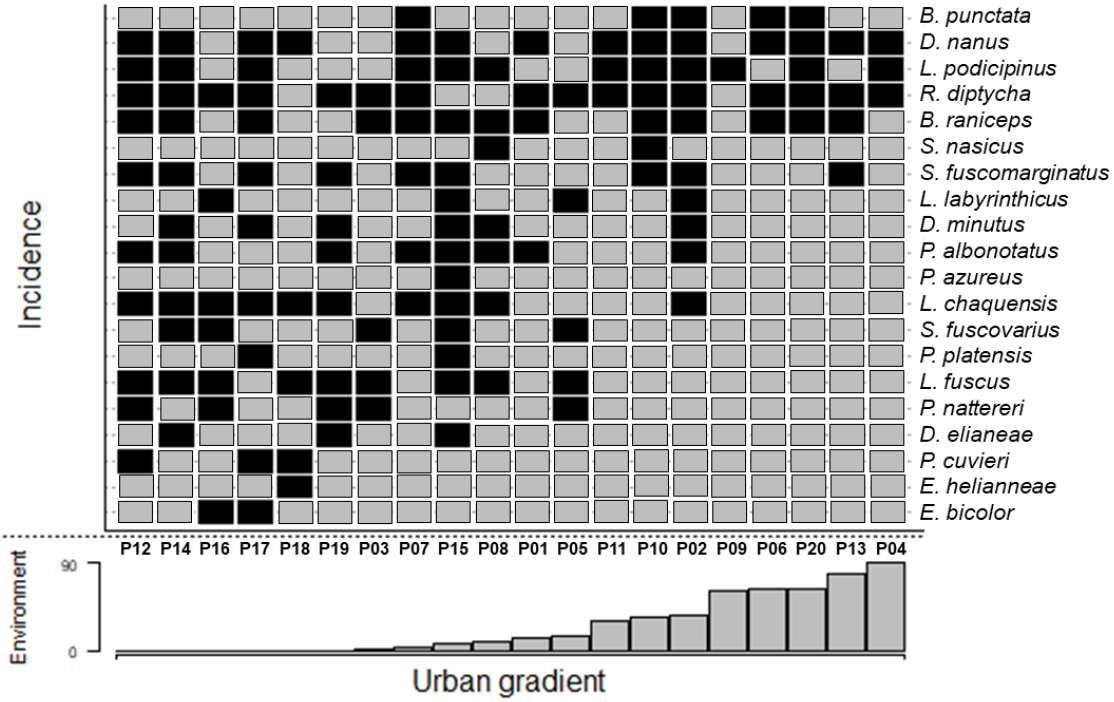
Incidence of all species ordered by the urbanization gradient (ranging from 0% to 89.4%, in a 500 m buffer). Black and gray blocks indicate the presence or absence of the species, respectively.

## 4. Discussion

Consistent with our predictions at the onset of this study, we found that anuran species richness was positively related to pond size and negatively associated with the degree of urbanization. In contrast to our initial predictions, total dissimilarity was mostly explained by turnover rather than nestedness, and turnover was best explained by the amount of urban infrastructure surrounding ponds.

The effect of pond size on species richness represents a positive species-area relationship (SARs), which is one of the most pervasive and robust patterns in ecology (Arrhenius 1921, Preston 1960, Connor and McCoy 2001), and has been described in urban context for a variety of organisms such as plants (Bräuniger et al. 2010), birds (Aida et al. 2016, Callaghan et al. 2018), and mammals (Matthies et al. 2017). Positive SARs are most commonly assumed to be caused by greater habitat heterogeneity in larger ponds supporting a larger number of species in larger areas (Williams 1943), as well as increased availability of resources allowing larger populations, which have low extinction probabilities (Hanski 1994). While these processes are likely to operate here, pond size might additionally be important in mitigating edge effects, with larger areas are relatively less affected by edge effects compared to smaller areas (given a similar shape) (Connor and McCoy 2001). Thereby, within an urban context, local disturbances at edges of ponds (Egger 2006, Du et al. 2010, Hu et al. 2018) might be less likely to affect the entire pond in large ponds, thereby decreasing the impacts of disturbance. In addition, a greater amount of water in larger ponds, may dilute the effects of pollution in larger ponds (Abel 1996).

We found evidence of a negative association between species richness and urban infrastructure. This result may be related to inhospitable characteristics of the urban environment for amphibians, such as artificial light (Baker and Richardson 2006), noise (Leon et al. 2019, Yi and Sheridan 2019), roads (Carr and Fahrig 2001, Pillsbury and Miller 2008), and possible contaminants (Knutson et al. 1999). Such drivers may influence the anuran life cycle in several ways. Noise pollution and artificial light can directly affect amphibian calling behavior, potentially leading to changes in survival and reproduction rates (Sun and Narins 2005, Baker and Richardson 2006). In addition, roads can increase mortality during dispersal and thereby increase probability of local extinction (Ashley and Robinson 1996, Findlay et al. 2001, Rubbo and Kiesecker 2005). Furthermore, since anurans feed on invertebrates, and several studies have found a decrease in invertebrate diversity as well as abundance in urban environments (e.g., Beavan et al. 2001, Moore and Palmer 2005, Jones and Leather 2013), urbanizations may reduce the availability of resources for anurans. A combination of these factors may negatively influence the ability of species to maintain viable populations in ponds in areas with a high degree of urbanization.

Contrary our predictions at the outset of this study, and the results from other studies (e.g., González-Oreja et al. 2012, Ficetola and De Bernardi 2004), we found that differences in community composition among ponds were caused primarily by turnover. However, similar results were found by Rádková et al. (2014) and Specziár et al. (2018) among invertebrate communities in aquatic environments in non-urban context. Our results suggest that urbanization is acting as an environmental filter (Piano et al. 2019), leading to the replacement of species not adapted to urban conditions by species that are more efficient in exploiting urban resources (McKinney 2006, Menke et al. 2011). As a result of such turnover of species, communities in ponds in highly urbanized areas are similar to each other, a process known as biotic homogenization (McKinney and Lockwood 1999, Olden and Rooney 2006). Biotic homogenization is a likely common consequence of anthropogenic disturbance (McKinney 2006, Olden 2006, Schwartz et al. 2006), and can have potential consequences for a variety of ecosystem services (Schwartz et al. 2006). Thereby, our results show that in more urbanized areas anuran communities are not only losing species, but also are becoming more homogeneous, which highlights a need for strengthened conservation efforts in the face of increasing replacement of natural habitat by urban areas (Grimm et al. 2008, Seto et al. 2011, 2012).

We found that the importance of different explanatory variables differed between species richness and community composition. For example, pond size was more important for the number rather than the identity of species. Similarly, forest cover was not important in models of species richness, but was a key predictor of the turnover component of beta diversity. This result highlights the need to protect forest habitat when the goal is to conserve habitat specialist species in peri-urban environments.

### Management implications

Our results provide valuable information for effective conservation of anurans in urban environments. While it is not feasible to reduce the percentage of urbanized areas to increase anuran richness, focusing conservation efforts on large ponds in highly urbanized environments might be an effective strategy to maintain a maximum number of species. Therefore, the negative effect of urbanization on amphibians could be reduced by increasing pond size. Although relationships among anuran communities and urbanization areas is far from being completely understood, and more studies focusing on anuran in urban context are necessary, our results may provide useful information for on-the-ground conservation actions.

## Acknowledgments

This study was financed in part by the Coordenação de Aperfeiçoamento de Pessoal de Nível Superior – Brasil (CAPES) – Finance Code 001 and Universidade Federal de Mato Grosso do Sul – UFMS/MEC – Brasil.

## Author contribution

CCG and MAG conceived the idea, designed the study and collect field data. CCG, MAG, and DBP performed the data analyses. All authors commented on drafts and gave final approval for publication.

## Notes

### Competing Interest Statement

The authors have declared no competing interest.

## References

Abel PD (1996) Water pollution problems and solutions. In: Water pollution biology. 2 ed. CRC Press. pp 1–27.

Aida N, Sasidhran S, Kamarudin N et al (2016) Woody trees, green space and park size improve avian biodiversity in urban landscapes of Peninsular Malaysia. Ecol Indic, 69, 176–183. https://doi.org/10.1016/j.ecolind.2016.04.025

Almeida-Gomes M, Rocha CF, Vieira MV (2016) Local and Landscape Factors Driving the Structure of Tropical Anuran Communities: Do Ephemeral Ponds have a Nested Pattern? Biotropica 48(3):365–372. https://doi.org/10.1111/btp.12285

Anderson MJ, Crist TO, Chase JM et al (2011) Navigating the multiple meanings of β diversity: a roadmap for the practicing ecologist. Ecol Lett 14:19–28. https://doi.org/10.1111/j.1461-0248.2010.01552.x

Arrhenius O (1921) Species and Area. J of Ecol 9:95–99.

Ashley EP, Robinson JT (1996) Road mortality of amphibians, reptiles and other wildlife on the Long Point Causeway, Lake Erie, Ontario. Can Field Nat 110(3):403–412.

Baker BJ, Richardson JML (2006) The effect of artificial light on male breeding-season behaviour in green frogs, *Rana clamitans melanota*. Can J of Zool 84(10): 1528–1532. https://doi.org/10.1139/z06-142

Barton K (2018) MuMIn: Multi-Model Inference. R package version 1.42.1. https://CRAN.R-project.org/package=MuMIn

Baselga A, Orme D, Villeger S, et al (2018) betapart: Partitioning Beta Diversity into Turnover and Nestedness Components. R package version 1.5.1. https://CRAN.R-project.org/package=betapart

Baselga A (2010) Partitioning the turnover and nestedness components of beta diversity. Glob Ecol and Biogeogr 19:134–143. https://doi.org/10.1111/j.1466-8238.2009.00490.x

Batáry P, Kurucz K, Suarez-Rubio M, Chamberlain DE (2018) Non-linearities in bird responses across urbanization gradients: A meta-analysis. Glob Chang Biol 24(3): 1046–1054. https://doi.org/10.1111/gcb.13964

Beavan L, Sadler J, Pinder C (2001) The invertebrate fauna of a physically modified urban river. Hydrobiol 445:97–108. https://doi.org/10.1023/A:1017584105641

Bender MG, Leprieur F, Mouillot D et al (2017) Isolation drives taxonomic and functional nestedness in tropical reef fish faunas. Ecography 40(3):425–435. https://doi.org/10.1111/ecog.02293

Bräuniger C, Knapp S, Kühn I, Klotz S (2010) Testing taxonomic and landscape surrogates for biodiversity in an urban setting. Landsc and Urban Plan 97(4):283–295. https://doi.org/10.1016/j.landurbplan.2010.07.001

Callaghan CT, Major RE, Lyons MB et al (2018) The effects of local and landscape habitat attributes on bird diversity in urban greenspaces. Ecosphere 9(7). https://doi.org/10.1002/ecs2.2347

Carr LW, Fahrig L (2001) Effect of road traffic on two amphibian species of differing vagility. Conserv Biol 15(4):1071–1078. https://doi.org/10.1046/j.1523-1739.2001.0150041071.x

Christie FJ, Cassis G, Hochuli DF (2010) Urbanization affects the trophic structure of arboreal arthropod communities. Urban Ecosyst 13(2):169–180. https://doi.org/10.1007/s11252-009-0115-x

Connor EF, McCoy ED (2001) Species-area relationships. Encycl of Biodivers 5:397–411.

Crump, ML, Scott NJ (1994) Visual encounter survey. In: Heyer, W.R. Donnelly, MA; McDiarmid, R.W, Donnelly, Heyek, L.C., and Foster, M.S. (Eds) Measuring and monitoring Biological diversity, Standard Methods for Amphibians Smithsonian Institution Press, Washington D.C: pp 84–91

Dale S (2018) Urban bird community composition influenced by size of urban green spaces, presence of native forest, and urbanization. Urban Ecosyst 21(1):1–14. https://doi.org/10.1007/s11252-017-0706-x

Davis TM, Ovaska K (2001) Individual recognition of amphibians: effects of toe clipping and fluorescent tagging on the salamander *Plethodon vehiculum*. J of Herpetol 35(2):217–225.

Dickman CR (1987) Habitat fragmentation and vertebrate species richness in an urban environment. J of Appl Ecol 24(2):337–351.

Doan TM (2003) Which methods are most effective for surveying rain forest herpetofauna?. J of Herpetol 37(1):72–81. https://doi.org/10.1670/0022-1511(2003)037[0072:WMAMEF]2.0.CO;2

Du N, Ottens H, Sliuzas R (2010) Spatial impact of urban expansion on surface water bodies - A case study of Wuhan, China. Landsc and Urban Plann 94(3-4):175–185. https://doi.org/10.1016/j.landurbplan.2009.10.002

Egger S (2006) Determining a sustainable city model. Env Modell & Softw 21(9):1235–1246. https://doi.org/10.1016/j.envsoft.2005.04.012

Fernández-Juricic E (2002) Can human disturbance promote nestedness? A case study with breeding birds in urban habitat fragments. Oecologia 131(2):269–278. https://doi.org/10.1007/s00442-002-0883-y

Ferreira CMM, de Aquino Ribas AC, de Souza FL (2017) Species composition and richness of anurans in Cerrado urban forests from central Brazil. Acta Herpetol 12(2):157–165.

Ficetola GF, De Bernardi F (2004) Amphibians in a human-dominated landscape: the community structure is related to habitat features and isolation. Biol Conserv 119(2):219–230.

Ficetola GF, Marziali L, Rossaro B, et al (2011) Landscape-stream interactions and habitat conservation for amphibians. Ecol Appl 21(4):1272–1282.

Findlay CS, Lenton J, Zheng L (2001) Land-use correlates of anuran community richness and composition in southeastern Ontario wetlands. Ecoscience 8(3):336–343. https://doi.org/10.1080/11956860.2001.11682661

Fischer JD, Cleeton SH, Lyons TP, Miller JR (2012) Urbanization and the predation paradox: the role of trophic dynamics in structuring vertebrate communities. Biosci 62(9):809–818. https://doi.org/10.1525/bio.2012.62.9.6

Flint WD, Harris RN (2005) The efficacy of visual encounter surveys for population monitoring of *Plethodon punctatus* (Caudata: Plethodontidae). J of Herpetol 39(4):578–584. https://doi.org/10.1670/255-04A.1

Gagné SA, Fahrig L (2007) Effect of landscape context on anuran communities in breeding ponds in the National Capital Region, Canada. Landsc Ecol 22(2):205–215. https://doi.org/10.1007/s10980-006-9012-3

González-Oreja JA, De La Fuente-Díaz AA, Hernández-Santín L et al (2012) Can human disturbance promote nestedness? Songbirds and noise in urban parks as a case study. Landsc and Urban Plann 104(1):9–18. https://doi.org/10.1016/j.landurbplan.2011.09.001

Grimm NB, Faeth SH, Golubiewski NE et al (2008) Global change and the ecology of cities. Science 319(5864):756–760

Grover MC (2006) Comparative effectiveness of nighttime visual encounter surveys and cover object searches in detecting salamanders. Herpetol Conserv and Biol 1(2):93–99.

Hanski I (1994). Patch-occupancy dynamics in fragmented landscapes. Trends in Ecol & Evol 9(4):131–135. https://doi.org/10.1016/0169-5347(94)90177-5

Hu A, Li S, Zhang L et al (2018) Prokaryotic footprints in urban water ecosystems: A case study of urban landscape ponds in a coastal city, China. Env Pollut 242:1729–1739. https://doi.org/10.1016/j.envpol.2018.07.097

IUCN (2020) The IUCN Red List of Threatened Species. Version 2020-2. https://www.iucnredlist.org. Accessed 15 May 2020.

Johansson F, Bini LM, Coiffard P et al (2019) Environmental variables drive differences in the beta diversity of dragonfly assemblages among urban stormwater ponds. Ecol Indic 106. https://doi.org/10.1016/j.ecolind.2019.105529

Johnson PT, Hoverman JT, McKenzie VJ et al (2013) Urbanization and wetland communities: applying metacommunity theory to understand the local and landscape effects. J of Appl Ecol 50(1):34–42. https://doi.org/10.1111/1365-2664.12022

Jones EL, Leather SR (2013) Invertebrates in urban areas: a review. Eur J Entomol 109(4):463–478.

Jupke JF, Schäfer RB (2020). Should ecologists prefer model-over distance-based multivariate methods?. Ecol and Evol 10(5):2417–2435. https://doi.org/10.1002/ece3.6059

Kinzig AP, Grove JM (2001) Urban-suburban ecology. In: Encyclopedia of Biodiversity (Eds. SA Levin). Academic Press, San Diego, pp 733–745

Knutson MG, Sauer JR, Olsen DA et al (1999) Effects of landscape composition and wetland fragmentation on frog and toad abundance and species richness in Iowa and Wisconsin, USA. Conserv Biol 13(6):1437–1446. https://doi.org/10.1046/j.1523-1739.1999.98445.x

Leibold MA, Chase JM (2018) Process in metacommunities. In: Metacommunity ecology, Princeton University Press, pp 49–88.

Leon E, Peltzer PM, Lorenzon R et al (2019) Effect of traffic noise on *Scinax nasicus* advertisement call (Amphibia, Anura). Iheringia. Ser Zool 109. http://dx.doi.org/10.1590/1678-4766e2019007

Lima NG, Oliveira U, Souza RC, Eterovick PC (2019) Dynamic and diverse amphibian assemblages: Can we differentiate natural processes from human induced changes?. PloS One 14(3). https://doi.org/10.1371/journal.pone.0214316

Lüdecke D (2019) _sjstats: Statistical Functions for Regression Models (Version 0.17.3). doi: 10.5281/zenodo.1284472 (URL: http://doi.org/10.5281/zenodo.1284472), <URL: https://CRAN.R-project.org/package=sjstats>.

Matthies SA, Rüter S, Schaarschmidt F, Prasse R (2017) Determinants of species richness within and across taxonomic groups in urban green spaces. Urban Ecosyst 20(4):897–909. https://doi.org/10.1007/s11252-017-0642-9

Maxwell SL, Fuller RA, Brooks TM, Watson JE (2016) Biodiversity: The ravages of guns, nets and bulldozers. Nat News 536(7615):143–145. https://doi.org/10.1038/536143a

McDonnell MJ, Hahs AK (2008) The use of gradient analysis studies in advancing our understanding of the ecology of urbanizing landscapes: current status and future directions. Landsc Ecol 23(10):1143–1155. https://doi.org/10.1007/s10980-008-9253-4

McDonnell MJ, Pickett STA, Groffman P et al (1997) Ecosystem processes along an urban-to-rural gradiente. Urban Ecosyst 1:21–36. https://doi.org/10.1023/A:1014359024275

McKinney ML (2006). Urbanization as a major cause of biotic homogenization. Biol Conserv 127(3):247–260. https://doi.org/10.1016/j.biocon.2005.09.005

McKinney ML (2008) Effects of urbanization on species richness: a review of plants and animals. Urban Ecosyst 11(2):161–176. https://doi.org/10.1007/s11252-007-0045-4

McKinney ML, Lockwood JL (1999) Biotic homogenization: a few winners replacing many losers in the next mass extinction. Trends in Ecol & Evol 14(11):450–453. https://doi.org/10.1016/S0169-5347(99)01679-1

Menke SB, Guénard B, Sexton JO et al (2011) Urban areas may serve as habitat and corridors for dry-adapted, heat tolerant species; an example from ants. Urban Ecosyst 14(2):135–163. https://doi.org/10.1007/s11252-010-0150-7

Miller JR, Hobbs RJ (2002) Conservation where people live and work. Conserv Biol 16(2):330–337.

Moore AA, Palmer MA (2005) Invertebrate biodiversity in agricultural and urban headwater streams: implications for conservation and management. Ecol Appl 15(4):1169–1177. https://doi.org/10.1890/04-1484

Nagelkerke NJD (1991) A note on a general definition of the coefficient of determination. Biom 78:691–692. doi.org/10.1093/biomet/78.3.691

Naimi B, Hamm NAS, Groen TA et al (2014) Where is positional uncertainty a problem for species distribution modelling?. Ecography 37:191–203. https://doi.org/10.1111/j.1600-0587.2013.00205.x

Oksanen J, Blanchet FG, Friendly M et al (2019) vegan: Community Ecology Package. R pac kage version 2.5-6. https://CRAN.R-project.org/package=vegan

Olden JD (2006) Biotic homogenization: a new research agenda for conservation biogeography. J of Biogeogr 33(12):2027–2039. https://doi.org/10.1111/j.1365-2699.2006.01572.x

Olden JD, Rooney TP (2006) On defining and quantifying biotic homogenization. Glob Ecol and Biogeogr 15(2): 113–120. https://doi.org/10.1111/j.1466-822X.2006.00214.x

Parris KM. 2006. Urban amphibian assemblages as metacommunities. J of Anim Ecol 75(3):757–764. https://doi.org/10.1111/j.1365-2656.2006.01096.x

Parris KM, Amati M, Bekessy SA et al (2018) The seven lamps of planning for biodiversity in the city. Cities 83:44–53. https://doi.org/10.1016/j.cities.2018.06.007

Piano E, Souffreau C, Merckx T et al (2019) Urbanization drives cross-taxon declines in abundance and diversity at multiple spatial scales. Glob Chang Biol 26(3): 1196–1211. https://doi.org/10.1111/gcb.14934

Pillsbury FC, Miller JR (2008) Habitat and landscape characteristics underlying anuran community structure along an urban–rural gradient. Ecol Appl 18(5): 1107–1118. https://doi.org/10.1890/07-1899.1

PLANURB – Agência Municipal de Meio Ambiente e Planejamento Urbano (2019). In: Perfil Socioeconômico Campo Grande, Mato Grosso do Sul. 26:41–76. http://www.capital.ms.gov.br/sisgran/#/. Accessed 05 November 2019.

Preston FW (1960) Time and space and the variation of species. Ecol 41(4):612–627.

Projeto MapBiomas – Coleção 4.0 da Série Anual de Mapas de Cobertura e Uso de Solo do Brasil. http://mapbiomas.org. Accessed 10 September 2019

R Core Team (2018) R: A language and environment for statistical computing. R Foundation for Statistical Computing, Vienna, Austria. URL: https://www.R-project.org/.

Rádková V, Bojková J, Křoupalová V et al (2014) The role of dispersal mode and habitat specialisation in metacommunity structuring of aquatic macroinvertebrates in isolated spring fens. Freshw Biol 59(11):2256–2267. https://doi.org/10.1111/fwb.12428

Ribeiro JF, Walter BMT (1998) Fitofisionomias do bioma Cerrado. In: Sano SM, Almeida Sp. (eds). Cerrado: ambiente e flora. Brasília, DF: Embrapa Cerrados, pp 89–166.

Rubbo MJ, Kiesecker JM (2005) Amphibian breeding distribution in an urbanized landscape. Conserv Biol 19(2):504–511. https://doi.org/10.1111/j.1523-1739.2005.000101.x

Sá RO, Tonini JFR, van Huss H et al (2018) Multiple connections between Amazonia and Atlantic Forest shaped the phylogenetic and morphological diversity of *Chiasmocleis Mehely*, 1904 (Anura: Microhylidae: Gastrophryninae). Mol Phylogenetics & Evol 130:198–210. https://doi.org/10.1016/j.ympev.2018.10.021

Schwartz MW, Thorne JH, Viers JH (2006) Biotic homogenization of the California flora in urban and urbanizing regions. Biol Conserv 127(3):282–291. https://doi.org/10.1016/j.biocon.2005.05.017

Semlitsch RD, Bodie JR (2003) Biological criteria for buffer zones around wetlands and riparian habitats for amphibians and reptiles. Conserv Biol 17(5):1219–1228. doi.org/10.1046/j.1523-1739.2003.02177.x

Seto KC, Fragkias M, Guneralp B et al (2011) A Meta-Analysis of Global Urban Land Expansion. PloS ONE 6(8):1–9. https://doi.org/10.1371/journal.pone.0023777

Seto KC, Guneralp B, Hutyra LR (2012) Global forecasts of urban expansion to 2030 and direct impacts on biodiversity and carbon pools. P Natl Acad Sci-Biol 109(40):16083–16088. https://doi.org/10.1073/pnas.1211658109

Slipinski P, Zmihorski M, Czechowski W (2012) Species diversity and nestedness of ant assemblages in an urban environment. Eur J of Entomol 109(2): 197–206.

Specziár A, Árva D, Tóth M et al (2018) Environmental and spatial drivers of beta diversity components of chironomid metacommunities in contrasting freshwater systems. Hydrobiol 819(1):123–143. https://doi.org/10.1007/s10750-018-3632-x

Sun JW, Narins PM (2005) Anthropogenic sounds differentially affect amphibian call rate. Biol Conserv 121(3):419–427. https://doi.org/10.1016/j.biocon.2004.05.017

United Nations (2018) World Urbanization Prospects, the 2018 Revision. Available at: http://esa.un.org/unpd/wup/index.htm. Accessed 31 July 2020

Wells KD (2007) The ecology and behavior of amphibians. University of Chicago Press, 1148 pp.

Whittaker RH (1960) Vegetation of the Siskiyou Mountains, Oregon and California. Ecol Monog 30:280–338.

Williams CB (1943) Area and number of species. Nat 152:264–267. https://doi.org/10.1038/152264a0

Yi YZ, Sheridan JA (2019) Effects of traffic noise on vocalisations of the rhacophorid tree frog *Kurixalus chaseni* (Anura: Rhacophoridae) in Borneo. Raffles Bull of Zool 67:77–82. https://doi.org/10.26107/rbz-2019-0007

Youngquist MB, Inoue K, Berg DJ, Boone MD (2017) Effects of land use on population presence and genetic structure of an amphibian in an agricultural landscape. Landsc Ecol 32(1):147–162. https://doi.org/10.1007/s10980-016-0438-y

Zhang W, Li B, Shu X et al (2016) Responses of anuran communities to rapid urban growth in Shanghai, China. Urban For & Urban Green 20:365–374. https://doi.org/10.1016/j.ufug.2016.10.005

