## Supplementar Material for "High species turnover shapes anuran community composition in ponds along an urban-rural gradient"

^1^ Pós-Graduação em Ecologia e Conservação, Universidade Federal de Mato Grosso do Sul, Campo Grande, Mato Grosso do Sul, 79002-970, Brazil.

^2^ Instituto de Biociências, Universidade Federal de Mato Grosso do Sul, Campo Grande, Mato Grosso do Sul, 79002-970, Brazil.

^3^ Göthenburg Global Biodiversity Centre, Göteborg, SE-450, Sweden.

^4^ Departamento de Ciências Ambientais, Universidade Federal de São Paulo - UNIFESP, 09913-030, Brazil.

^5^ Fenner School of Environment and Societ, Australian National University, Canberra, ACT, Australia.

Carolina Ganci orcid: 0000-0001-7594-8056

Diogo B. Provete orcid: 0000-0002-0097-0651

Thomas Püttker orcid: [0000-0003-0605-1442](https://orcid.org/0000-0003-0605-1442)

Mauricio Almeida-Gomes orcid: 0000-0001-7938-354X

David Lindenmayer orcid: 0000-0002-4766-4088

**Table S1** Pond size (Area; m²), percentage of native forest in a buffer of 500 (For_500) and 1000 m (For_1000), and percentage of urban infrastructure in a buffer of 500 (Ubr_500) and 1000 m (Ubr_1000) of the ponds used in this study

|  | Area (m²) | For_500 | For_1000 | Urb_500 | Urb_1000 |
| --- | --- | --- | --- | --- | --- |
| P1 | 610 | 29.45 | 26.51 | 12.90 | 7.37 |
| P2 | 2069 | 22.35 | 12.10 | 35.35 | 52.90 |
| P3 | 109 | 35.43 | 31.31 | 1.09 | 4.20 |
| P4 | 437 | 0.57 | 1.88 | 89.42 | 83.91 |
| P5 | 397 | 9.43 | 24.28 | 14.11 | 21.11 |
| P6 | 1416 | 16.15 | 9.94 | 62.90 | 76.56 |
| P7 | 957 | 34.38 | 20.01 | 3.35 | 13.40 |
| P8 | 604 | 11.58 | 11.84 | 7.85 | 14.22 |
| P9 | 167 | 15.97 | 11.65 | 60.72 | 68.61 |
| P10 | 341 | 35.72 | 19.41 | 33.38 | 43.40 |
| P11 | 224 | 18.73 | 16.48 | 30.40 | 46.88 |
| P12 | 448 | 15.44 | 16.33 | 0.00 | 0.24 |
| P13 | 539 | 14.98 | 6.83 | 77.95 | 84.54 |
| P14 | 778 | 30.07 | 24.95 | 0.00 | 0.00 |
| P15 | 3080 | 0.30 | 10.43 | 7.09 | 10.14 |
| P16 | 1009 | 6.19 | 12.16 | 0.00 | 7.73 |
| P17 | 1578 | 33.58 | 47.35 | 0.00 | 0.00 |
| P18 | 434 | 36.11 | 23.08 | 0.00 | 0.00 |
| P19 | 287 | 19.42 | 17.47 | 0.00 | 0.00 |
| P20 | 488 | 20.74 | 19.41 | 63.85 | 66.75 |

**Table S2** Abundance of anuran species sampled at the 20 ponds surveyed.

|  | **P1** | **P2** | **P3** | **P4** | **P5** | **P6** | **P7** | **P8** | **P9** | **P10** | **P11** | **P12** | **P13** | **P14** | **P15** | **P16** | **P17** | **P18** | **P19** | **P20** |
| --- | --- | --- | --- | --- | --- | --- | --- | --- | --- | --- | --- | --- | --- | --- | --- | --- | --- | --- | --- | --- |
| **Bufonidae** |  |  |  |  |  |  |  |  |  |  |  |  |  |  |  |  |  |  |  |  |
| *Rhinella diptycha* | 1 | 2 | 2 | 1 | 2 | 1 | 2 | 0 | 0 | 1 | 13 | 5 | 2 | 1 | 0 | 12 | 1 | 0 | 1 | 8 |
| **Hylidae** |  |  |  |  |  |  |  |  |  |  |  |  |  |  |  |  |  |  |  |  |
| *Boana punctata* | 0 | 32 | 0 | 0 | 0 | 5 | 11 | 0 | 0 | 4 | 0 | 0 | 0 | 0 | 0 | 0 | 0 | 0 | 0 | 8 |
| *Boana raniceps* | 4 | 13 | 1 | 0 | 0 | 3 | 3 | 4 | 0 | 2 | 0 | 8 | 7 | 9 | 3 | 0 | 2 | 0 | 0 | 3 |
| *Dendropsophus elianeae* | 0 | 0 | 0 | 0 | 0 | 0 | 0 | 0 | 0 | 0 | 0 | 0 | 0 | 8 | 6 | 0 | 0 | 0 | 2 | 0 |
| *Dendropsophus minutus* | 0 | 18 | 0 | 0 | 0 | 0 | 0 | 1 | 0 | 0 | 0 | 0 | 0 | 36 | 15 | 0 | 5 | 0 | 1 | 0 |
| *Dendropsophus nanus* | 18 | 42 | 0 | 2 | 0 | 54 | 30 | 0 | 0 | 23 | 6 | 23 | 16 | 25 | 12 | 0 | 4 | 1 | 0 | 10 |
| *Pseudis platensis* | 0 | 0 | 0 | 0 | 0 | 0 | 0 | 0 | 0 | 0 | 0 | 0 | 0 | 0 | 7 | 0 | 7 | 0 | 0 | 0 |
| *Scinax fuscomarginatus* | 0 | 23 | 0 | 0 | 0 | 0 | 4 | 0 | 0 | 5 | 0 | 7 | 13 | 15 | 2 | 0 | 15 | 0 | 3 | 0 |
| *Scinax fuscovarius* | 0 | 0 | 1 | 0 | 6 | 0 | 0 | 0 | 0 | 0 | 0 | 0 | 0 | 3 | 2 | 3 | 0 | 0 | 0 | 0 |
| *Scinax nasicus* | 0 | 0 | 0 | 0 | 0 | 0 | 0 | 1 | 0 | 1 | 0 | 0 | 0 | 0 | 0 | 0 | 0 | 0 | 0 | 0 |
| **Leptodactylidae** |  |  |  |  |  |  |  |  |  |  |  |  |  |  |  |  |  |  |  |  |
| *Leptodactylus chaquensis* | 0 | 1 | 0 | 0 | 0 | 0 | 1 | 3 | 0 | 0 | 0 | 2 | 0 | 7 | 9 | 3 | 2 | 11 | 3 | 0 |
| *Leptodactylus fuscus* | 0 | 0 | 19 | 0 | 4 | 0 | 0 | 1 | 0 | 0 | 0 | 3 | 0 | 3 | 1 | 2 | 0 | 4 | 5 | 0 |
| *Leptodactylus labyrinthicus* | 0 | 4 | 0 | 0 | 2 | 0 | 0 | 0 | 0 | 0 | 0 | 0 | 0 | 0 | 2 | 1 | 0 | 0 | 0 | 0 |
| *Leptodactylus podicipinus* | 0 | 3 | 0 | 3 | 0 | 0 | 3 | 1 | 21 | 3 | 14 | 2 | 0 | 5 | 2 | 0 | 2 | 0 | 0 | 3 |
| *Physalaemus albonotatus* | 1 | 1 | 0 | 0 | 0 | 0 | 1 | 1 | 0 | 0 | 0 | 4 | 0 | 1 | 1 | 0 | 0 | 0 | 3 | 0 |
| *Physalaemus cuvieri* | 0 | 0 | 0 | 0 | 0 | 0 | 0 | 0 | 0 | 0 | 0 | 1 | 0 | 0 | 0 | 0 | 2 | 1 | 0 | 0 |
| *Physalaemus nattereri* | 0 | 0 | 1 | 0 | 1 | 0 | 0 | 0 | 0 | 0 | 0 | 1 | 0 | 0 | 0 | 1 | 0 | 0 | 1 | 0 |
| **Microhylidae** |  |  |  |  |  |  |  |  |  |  |  |  |  |  |  |  |  |  |  |  |
| *Elachistocleis bicolor* | 0 | 0 | 0 | 0 | 0 | 0 | 0 | 0 | 0 | 0 | 0 | 0 | 0 | 0 | 0 | 1 | 1 | 0 | 0 | 0 |
| *Elachistocleis helianneae* | 0 | 0 | 0 | 0 | 0 | 0 | 0 | 0 | 0 | 0 | 0 | 0 | 0 | 0 | 0 | 0 | 0 | 1 | 0 | 0 |
| **Phyllomedusidae** |  |  |  |  |  |  |  |  |  |  |  |  |  |  |  |  |  |  |  |  |
| *Pithecopus azureus* | 0 | 0 | 0 | 0 | 0 | 0 | 0 | 0 | 0 | 0 | 0 | 0 | 0 | 0 | 2 | 0 | 0 | 0 | 0 | 0 |
| **Total abundance** | **24** | **139** | **24** | **6** | **15** | **63** | **55** | **12** | **21** | **39** | **33** | **56** | **38** | **113** | **64** | **23** | **41** | **18** | **19** | **32** |

**Table S3** Observed and estimated richness by Chao 1 and mean sampling coverage (proportion of observed richness relative to richness estimated by Chao 1) per pond.

|  | Observed richness | Estimated richness | Mean sampling coverage |
| --- | --- | --- | --- |
| P1 | 4 | 5 | 0.80 |
| P2 | 10 | 10.5 | 0.95 |
| P3 | 5 | 6.5 | 0.76 |
| P4 | 3 | 3 | 1.00 |
| P5 | 5 | 5 | 1.00 |
| P6 | 4 | 4 | 1.00 |
| P7 | 8 | 8.5 | 0.94 |
| P8 | 7 | 17 | 0.41 |
| P9 | 1 | 1 | 1.00 |
| P10 | 7 | 7.5 | 0.93 |
| P11 | 3 | 3 | 1.00 |
| P12 | 10 | 10.3 | 0.97 |
| P13 | 4 | 4 | 1.00 |
| P14 | 11 | 12 | 0.91 |
| P15 | 13 | 13.1 | 0.99 |
| P16 | 7 | 8.5 | 0.82 |
| P17 | 10 | 10.2 | 0.98 |
| P18 | 5 | 8 | 0.62 |
| P19 | 8 | 9.5 | 0.84 |
| P20 | 5 | 5 | 1 |

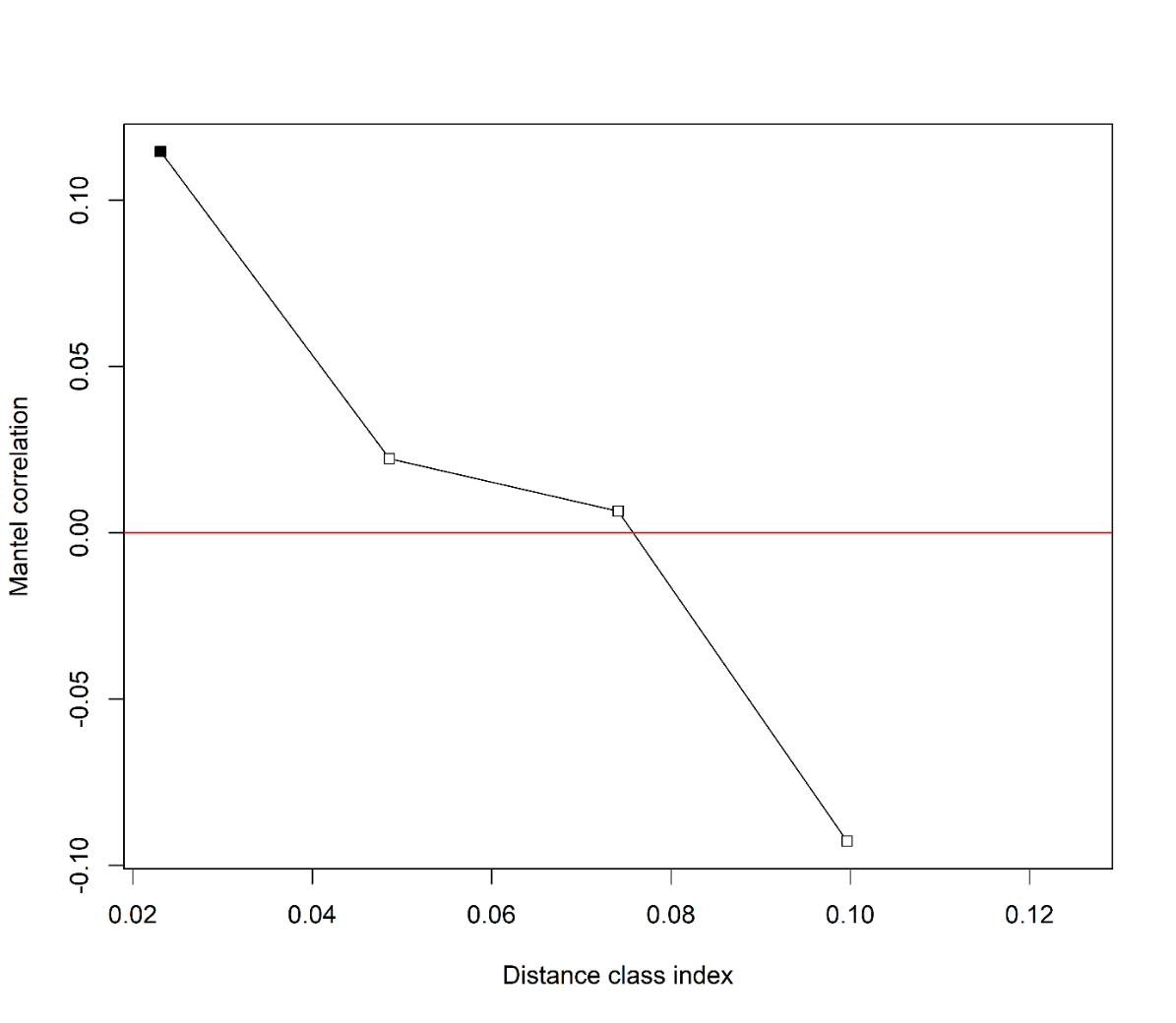

**Fig S1** Mantel spatial correlogram showing species dissimilarity along distance classes. Significant values are represented by black block, and non-significant ones are represented by white blocks. Notice there is only a slightly positive autocorrelation in the first distance class, that is, ponds separated by up to 300 m have more similar species composition than expected.

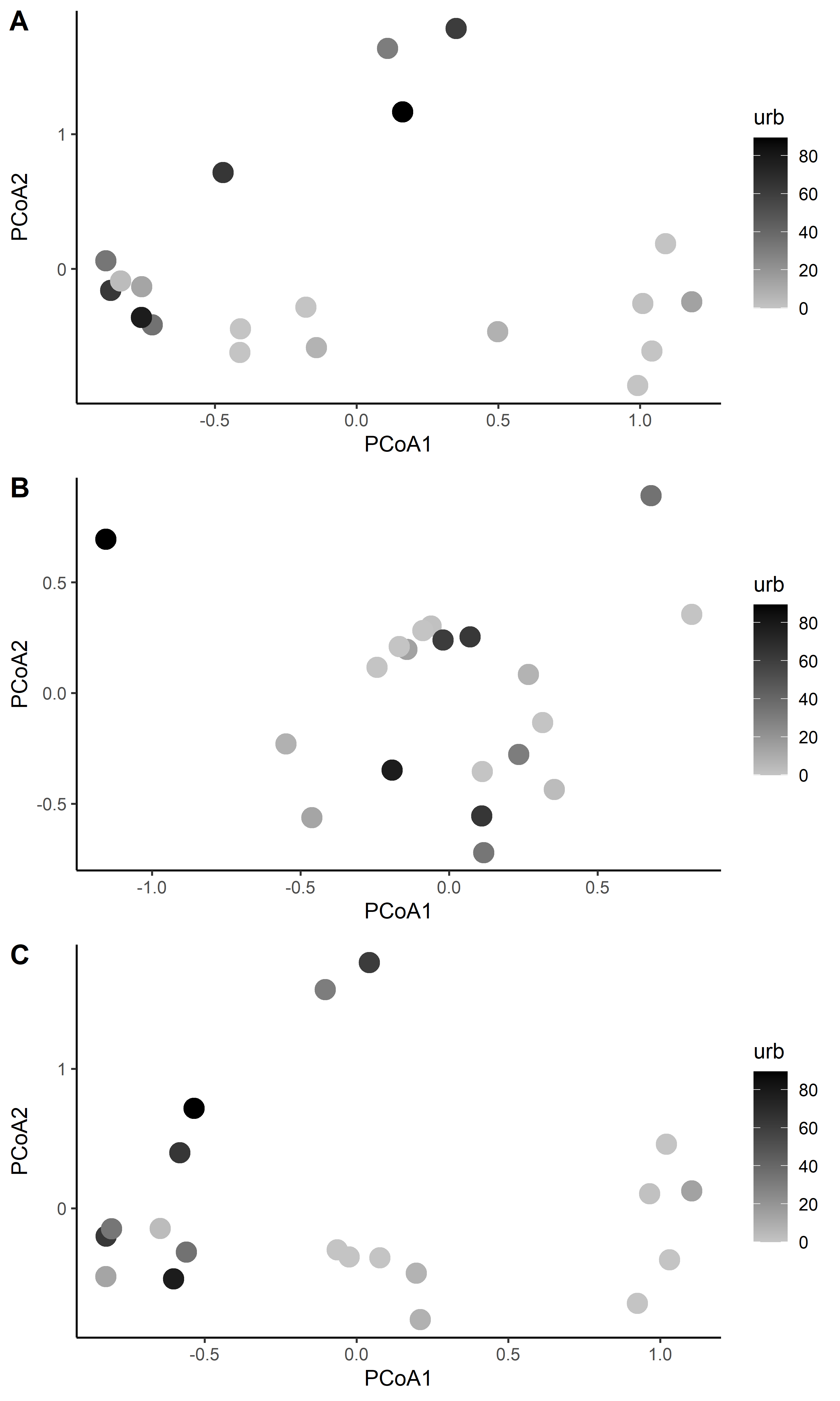

**Fig S2** Principal Coordinate Analysis showing the total taxonomic pair-wise dissimilarity between sites (A) and the relative importance of nestedness (B) and turnover (C), based on a distance matrix generated from the abundance data. Symbols are shaded according to the amount of urban infrastructure in a buffer of 500 m surrounding the central point of each pond (Urb_500).

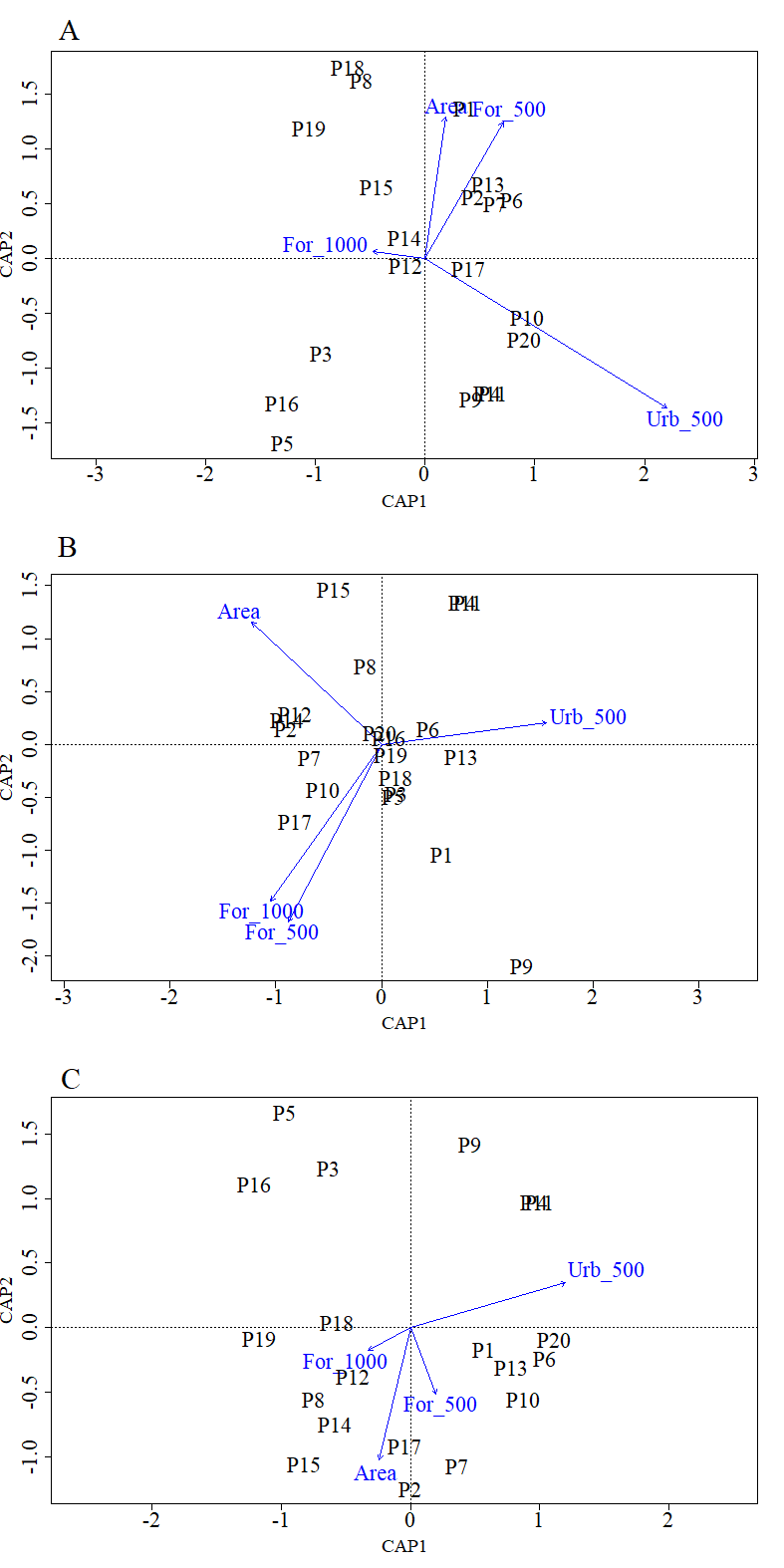

**Fig S3** Ordination diagram showing the result of the distance-based redundancy analysis (db-RDA) for (A) turnover component (B) nestedness component and, (C) total beta diversity, displaying the sites and landscape variables. Area – pond size. For_500 and For_100 – percentage of native forest in a buffer of 500 and 1,000 m respectively. Urb_500 – percentage of urban infrastructure in a buffer of 500 m.
